# Complete Mitochondrial and Plastid DNA Sequences of the Freshwater Green Microalga *Medakamo hakoo*

**DOI:** 10.1101/2021.07.27.453968

**Authors:** Mari Takusagawa, Shoichi Kato, Sachihiro Matsunaga, Shinichiro Maruyama, Yayoi Tsujimoto-Inui, Hisayoshi Nozaki, Fumi Yagisawa, Mio Ohnuma, Haruko Kuroiwa, Tsuneyoshi Kuroiwa, Osami Misumi

**Author notes:** Address correspondence to Osami Misumi.

## Abstract

Here we report the complete organellar genome sequences of *Medakamo hakoo*, a green alga identified in freshwater in Japan. It has 90.8-kb plastid and 36.5-kb mitochondrial genomes containing 80 and 33 putative protein coding genes, respectively, representing the smallest organellar genome among currently known core Trebouxiophyceae.

## Body

The genomes of various organisms have been sequenced, and knowledge has been accumulated about not only biodiversity but also universal genes that are conserved among all organisms. These findings will be updated as new species are discovered; therefore, exploring new species will continue to be important. Recently, we discovered a green alga, *Medakamo hakoo*, in freshwater in Japan (1, 2). Based on the preliminary microphotometric observation, we predicted that it would have the smallest organellar and nuclear genomes among green plants, which prompted us to investigate its genomes. Whole-genome sequencing was performed with the PacBio RS II system (Pacific Biosciences, Menlo Park, CA, USA), and the read sequences were assembled *de novo* with the RS HGAP Assemble.3 program run by SMRT Analysis software 2.3.0 (Pacific Biosciences). Genes encoded by the chloroplast/mitochondrial genomes were annotated using GeSeq (3) with the following organellar genomes as references: *Chlorella vulgaris* (NC_001865/NC_045362), *Coccomyxa subellipsoidea* C-169 (NC_015084/NC_015316), and *Botryococcus braunii* (NC_025545/NC_027722) because *M. hakoo* belongs to Trebouxiophyceae, as described in Kato *et al.* (submitted). StructRNAfinder (4) and RNAweasel (5) were used to predict rRNA. We obtained two contigs representing the organellar genomes (Fig.1). The chloroplast genome (cpDNA) is 90,872 bp and has a GC content of 41.4%. We functionally annotated 80 protein-coding genes, including 31 photosynthesis-related genes (components of photosystem I and II, the cytochrome *b*_6_*f* complex, and ATP synthase subunits) and 29 tRNA and 2 rRNA genes. The cpDNA of *M. hakoo* is slightly smaller than that of its most closely related alga, *Choricystis parasitica* (6); they share most genes but not their order. The mitochondrial genome (mtDNA) of *M. hakoo* is 36,544 bp and has a GC content of 35.2%. It is the smallest in size and the gene density appears high compared with other Trebouxiophyceae mtDNAs. It comprises 25 tRNAs, 3 rRNAs, and 33 predicted protein-coding genes, including 18 respiratory chain complex genes and ATP synthase, as well as *tatC* and a homing endonuclease gene.

**Fig. 1.**
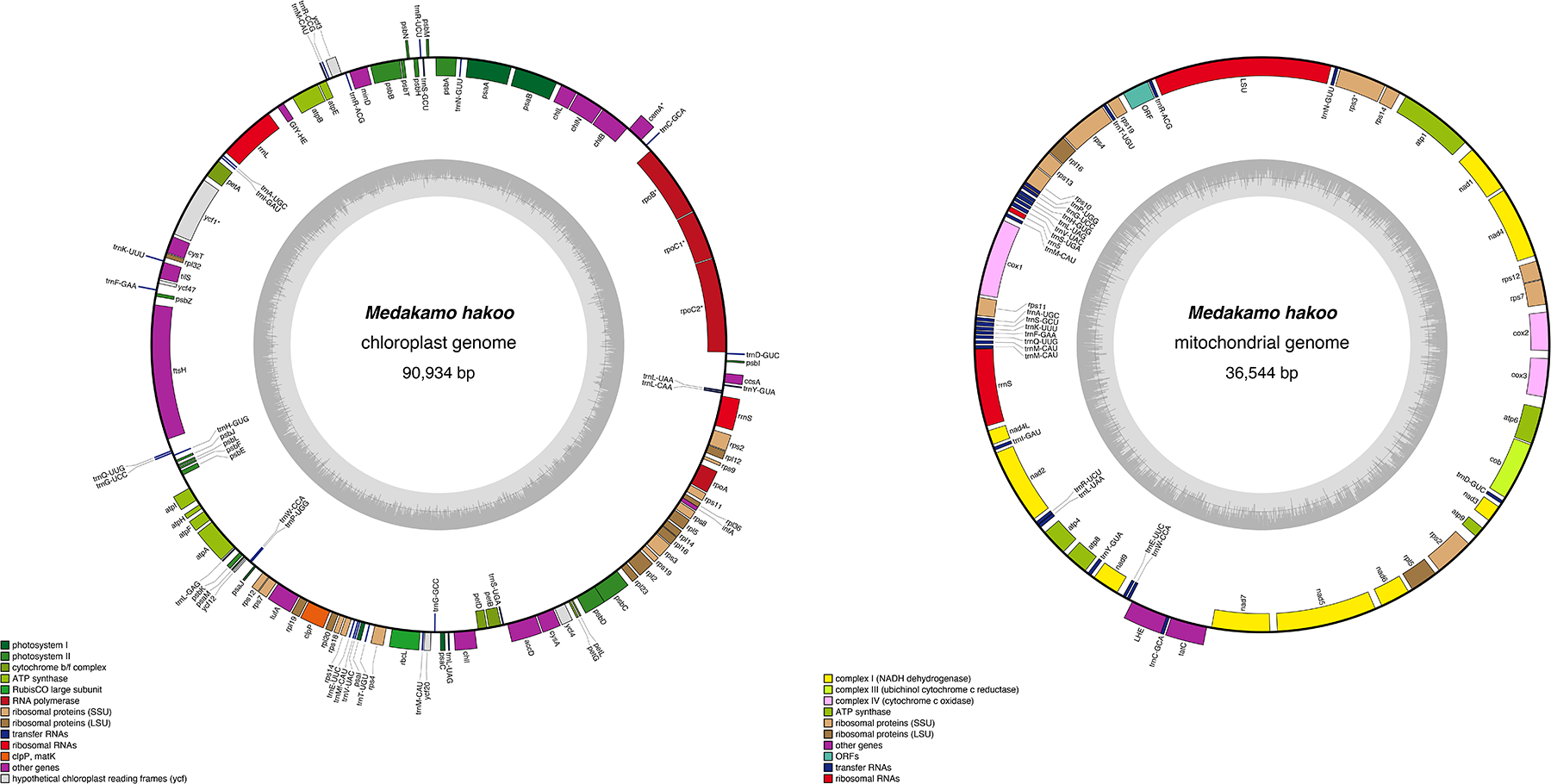
Complete chloroplast and mitochondrial genomes of *Medakamo hakoo*. The both genomes were plotted using OrganellarGenomeDRAW (11). The inner gray circle illustrates the G+C content of the genome.

Another notable characteristic of the mtDNA was that almost all the genes were localized to a single strand. This mtDNA characteristic is also observed among core Trebouxiales, including Trebouxiophyceae sp. MX-AZ01 (7) and *Coccomyxa subellipsoidea* C-169 (8) and *Botoryococcus braunii* (9,10); however, the group II introns of the *trnH* (gug), and *trnW* (cca) genes conserved in these species (10) were not predicted in *M. hakoo*. Normally in Trebouxiophyceae, *rpl10* gene was located next to *rps19* (10), but *M. hakoo* had another ORF that was not similar to other known genes. As expected, the size of the *M. hakoo* organellar genome was relatively small, and the gene density was high. It will be important for future algal research to focus on how mitochondrial genes encoded in a single strand are expressed (whether they form operons), and conversely how the separately encoded chloroplast rRNA genes, which should normally form an operon structure, exhibit coordinated transcription.

### Public availability and accession numbers

The cpDNA and mtDNA genome sequences were deposited in DNA Data Bank of Japan (https://www.ddbj.nig.ac.jp/index-e.html) with accession numbers LC604816 and LC604817, respectively.

## Acknowledgments

This work was supported by grants from the Ministry of Education, Culture, Sports, Science and Technology of Japan (16H04813 and 19H03260) to T.K. and by JST-CREST (Grant Number JPMJCR20S6) to S.Matsunaga. The authors thank Mr. Brody Frink (Kyoto University) for editing a draft of this manuscript.

## References

1. Kuroiwa T, Ohnuma M, Nozaki H, Imoto Y, Misumi O, Kuroiwa H. 2015. Cytological Evidence of Cell-Nuclear Genome Size of a New Ultra-Small Unicellular Freshwater Green Alga, “*Medakamo hakoo*” strain M-hakoo 311 I. Comparison with *Cyanidioschyzon merolae* and *Ostreococcus tauri*. Cytologia 80:143–150

2. Kuroiwa T, Ohnuma M, Imoto Y, Misumi O, Nagata N, Miyakawa I, Fujishima M, Yagisawa F, Kuroiwa H. 2016. Genome Size of the Ultrasmall Unicellular Freshwater Green Alga, Medakamo hakoo 311, as Determined by Staining with 4′,6-Diamidino-2-phenylindole after Microwave Oven Treatments: II. Comparison with *Cyanidioschyzon merolae*, *Saccharomyces cerevisiae* (n, 2n), and *Chlorella variabilis*. Cytologia 81:69–76

3. Tillich M, Lehwark P, Pellizzer T, Ulbricht-Jones ES, Fischer A, Bock R, Greiner S 2017. GeSeq – versatile and accurate annotation of organelle genomes. Nucleic Acids Research 45: W6–W11

4. Arias-Carrasco R, Vásquez-Morán, Y, Nakaya HI, Maracaja-Coutinho V. 2018. StructRNAfinder: an automated pipeline and web server for RNA families prediction. BMC Bioinformatics, 19: 55.

5. Lang BF, Laforest MJ, Burger G. 2007. Mitochondrial introns: a critical view. Trends Genet. 23:119–25

6. Lemieux C, Otis C, Turmel M. 2014. Chloroplast phylogenomic analysis resolves deep-level relationships within the green algal class *Trebouxiophyceae*. BMC Evol Biol. 14:211

7. Servín-Garcidueñas LE, Martínez-Romero E. 2012. Complete Mitochondrial and Plastid Genomes of the Green Microalga *Trebouxiophyceae* sp. Strain MX-AZ01 Isolated from a Highly Acidic Geothermal Lake. Eukaryotic Cell, 11:1417–1418

8. Smith DR, Burki F, Yamada T, Grimwood J, Grigoriev IV, et al. 2011. The GC-Rich Mitochondrial and Plastid Genomes of the Green Alga *Coccomyxa* Give Insight into the Evolution of Organelle DNA Nucleotide Landscape. PLOS ONE 6(8): e23624

9. Blifernez-Klassen O, Wibberg D, Winkler A, Blom J, Goesmann A, Kalinowski J, Kruse O. 2016. Complete chloroplast and mitochondrial genome sequences of the hydrocarbon oil-producing green microalga *Botryococcus braunii* Race B (Showa). Genome Announc. 4: e00524–16

10. Martínez-Alberola F, Barreno E, Casano LM, Gasulla F, Molins A, del Campo EM. 2019. Dynamic evolution of mitochondrial genomes in Trebouxiophyceae, including the first completely assembled mtDNA from a lichen-symbiont microalga (*Trebouxia* sp. TR9). Scientific Reports 9:8209

11. Greiner S, Lehwark P, Bock R. 2019. OrganellarGenomeDRAW (OGDRAW) version 1.3.1: expanded toolkit for the graphical visualization of organellar genomes. Nucleic Acids Research 47: W59–W64

